# The archaic CUP pilus SMF-1 utilizes a specific antiparallel bundling mechanism to initiate biofilm formation

**DOI:** 10.64898/2026.02.19.706768

**Authors:** Jessie Lynda Fields, Caitlyn C. Sebastian, Hua Zhang, Ayisha Zia, Abigail N. Robertson, Hui Wu, Megan R. Kiedrowski, Fengbin Wang

## Abstract

Bacterial biofilms represent a survival strategy that enables microbial communities to withstand environmental stress. *Stenotrophomonas maltophilia* is an emerging, antibiotic-resistant, Gram-negative, opportunistic pathogen that frequently colonizes the lungs of individuals with cystic fibrosis. Its chaperone-usher pathway (CUP) pilus, SMF-1, is present in nearly all clinical isolates and is essential for biofilm development; however, its molecular architecture has remained unknown. Here, we present a 4.0 Å cryo-EM structure revealing that SMF-1 is an archaic, rather than a classic, CUP pilus. SMF-1 forms thin, zigzag filaments that assemble into defined antiparallel “pili-couples,” which further aggregate into thick bundles. These bundles act as robust intercellular tethers, facilitating the rapid cell-to-cell aggregation required for biofilm initiation. Despite high sequence and structural similarity to classic CUP systems, SMF-1 lacks the interfaces required to form a rod-like architecture, suggesting it may represent an evolutionary intermediate between the CUP classes. Finally, we demonstrate that SMF-1 producing bacteria initiate biofilm formation within 24 hours and that flagella can further accelerate this process. Together, these findings uncover a conserved bundling mechanism that promotes bacterial colonization and may contribute to pathogenicity.

## INTRODUCTION

The formation of bacterial biofilms represents a sophisticated survival strategy that enables microbial communities to withstand diverse environmental stressors^1–4^. For pathogenic species, this structured lifestyle creates a formidable barrier that hinders the penetration of host immune cells and limits their defensive activity^5^. Beyond immune evasion, the biofilm architecture also confers tolerance to antibiotics, often requiring minimum inhibitory concentrations substantially higher than those needed for planktonic cells^6–9^. This inherent resilience frequently contributes to chronic infections and presents a major challenge for clinical treatment^10^.

Biofilm formation is largely driven by extracellularly secreted components, primarily the extracellular polymeric substance (EPS) matrix. This matrix, composed of functional protein appendages, polysaccharides, and extracellular DNA, provides the physical architecture necessary for community stability and protection against external stress^11–13^. Among the protein appendages, several major classes of filaments have been linked to biofilm development. Bacterial flagella, which enable swimming and swarming motility, are integral to early stages of surface attachment and biofilm initiation; however, they are frequently downregulated as biofilms mature^14,15^. Bacterial pili, including type IV pili (T4P) and chaperone-usher pathway (CUP) pili, have also been implicated in biofilm formation through distinct mechanisms. Type IV pili facilitate biofilm formation by mediating initial attachment to surfaces, enabling twitching motility, and promoting cell-to-cell aggregation^16–18^. In contrast, the role of CUP pili is more complex, reflecting the diversity of their various subgroups.

CUP pili are among the most abundant bacterial pili in Gram-negative species. Related pili structures are also present in Gram-positive bacteria^19^ and archaea^20,21^, although their corresponding assembly mechanisms remain unknown. Based on the outer membrane usher sequence and pilus operon organization, CUP pili are subdivided into three classes: classical, alternate, and archaic. Both classical^22,23^ and alternate^24^ CUP pili form rod-like structures, and their contribution to biofilm formation, when present, is likely mediated primarily through initial surface attachment. Archaic CUP pili, by contrast, form thinner, zigzag^25,26^ structures as single filaments. These single pili are often observed to form thick bundles bridging different cells. If these pili pack in an antiparallel fashion within the bundles, the filaments likely originate from different cells. Therefore, bundling could provide a robust mechanism for sustaining cell-to-cell aggregation and driving early biofilm formation. Many pathogens utilize CUP pili to establish and maintain infection, and their role in virulence can be crucial. For example, the emerging opportunistic pathogen *Stenotrophomonas maltophilia* frequently colonizes damaged, muco-obstructed airways in people with cystic fibrosis and often appears alongside or following *Pseudomonas aeruginosa* infections^27^. Beyond respiratory colonization, *S. maltophilia* has also been reported to promote tumor progression by triggering degradation of STING (stimulator of interferon genes), thereby dampening host’s anti-cancer immune responses^28^. Central to its pathogenicity is the SMF-1 pilus, a CUP system that is not only highly conserved at the sequence level and found in nearly all *S. maltophilia* clinical isolates^29–34^. Previously annotated as a classical CUP pilus^35^ (γ-clade), SMF-1 is essential for initial attachment^35^ and biofilm maturation on both abiotic and biotic surfaces^36,37^. Rather than functioning solely as a simple adhesin typical of many classic CUP pili, recent studies indicate that SMF-1 pili can act as physical tethers that facilitate rapid cell-to-cell aggregation and interspecies interactions, forming a defensive shield that enhances survival^38^.

In this work, we determined the cryo-EM structure of the SMF-1 pilus. Surprisingly, we found that SMF-1 is an archaic CUP pilus consisting of single fibers with a characteristic zigzag morphology that assemble into thick bundles. Using asymmetrical reconstruction, we reveal the antiparallel nature of the pilus arrangement within these bundles. Two SMF-1 pili form an antiparallel “pilus-couple,” and this unit further aggregates into larger bundles. Furthermore, we show that although SMF-1 resembles classical CUP pili at the levels of sequence, overall structure, and pilin-cluster organization, these pilins lack the interface required to form key interactions that underlie classical CUP pilus architecture. Finally, we investigated the biofilm initiation capability of *S. maltophilia* while monitoring appendage production, establishing that SMF-1 production is dynamic and can drive biofilm initiation independently. Moreover, we show that this process can be further accelerated in the presence of bacterial flagellar filaments.

## RESULTS

### Cryo-EM structure of the SMF-1 pilus reveals a specific antiparallel bundling mechanism

It has been previously reported in various studies that *Stenotrophomonas maltophilia* SMF-1 pili are essential for initial attachment and subsequent biofilm formation on both abiotic and biotic surfaces^32,34–36,39^. In our previous preparations of flagellar filaments^40^ from *S. maltophilia*, we observed large amounts of pili bundles, with single fibers measuring approximately 5 nm in diameter. When we culture *S. maltophilia* in liquid media, these types of pili bundles, presumably SMF-1 pili, are frequently observed between cells (Fig. 1A), consistent with previous TEM (transmission electron microscope) observations^36^.

**Figure 1.**
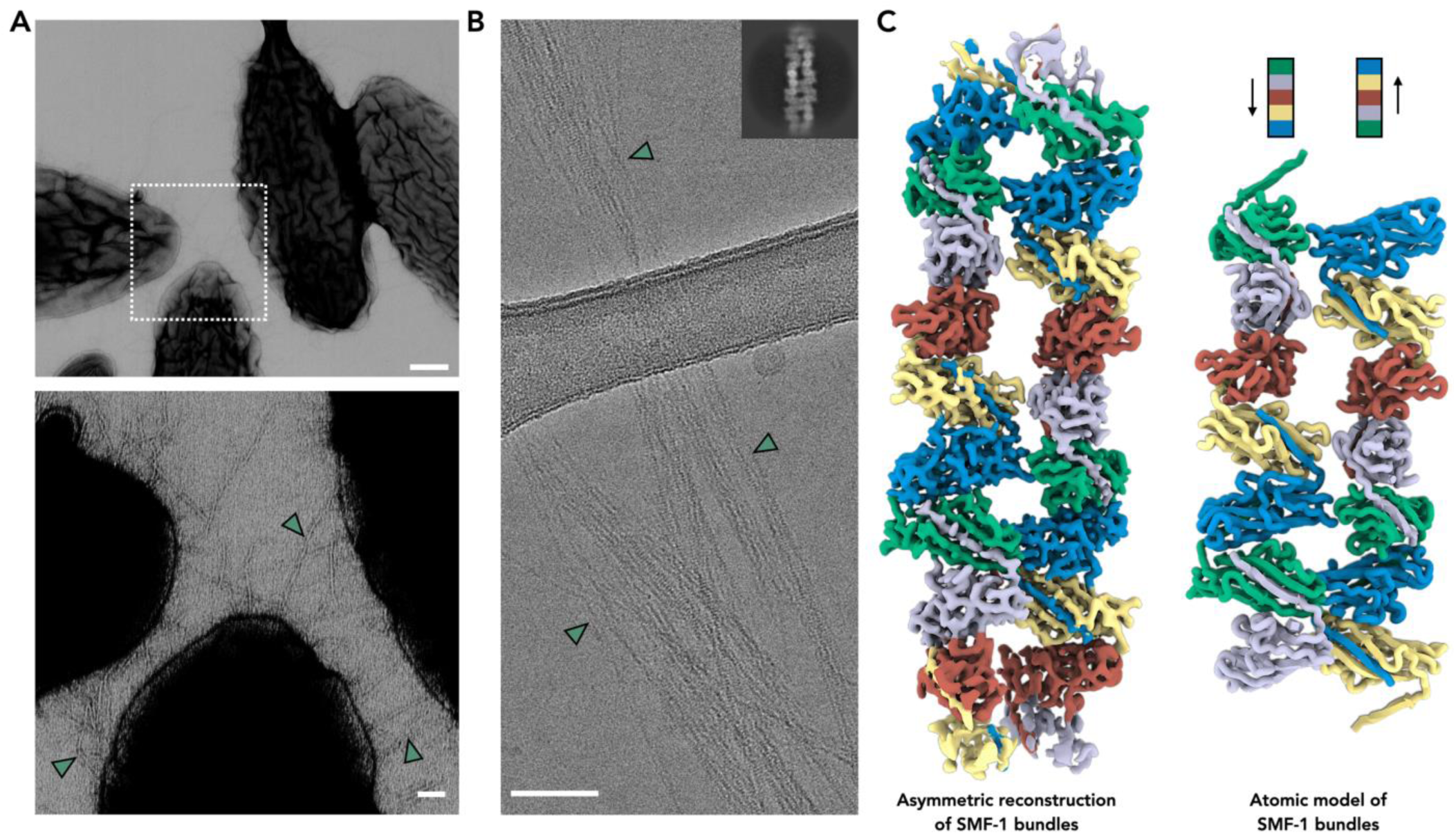
Cryo-EM of the SMF-1 pilus from *S. maltophilia*. (A) Negative staining image of *S. maltophilia* cells (top). The area between three cells is further magnified in the bottom panel. Pili bundles located between cells are indicated with green arrowheads. Scale bar for the top negative staining image, 200 nm. Scale bar for the bottom zoomed-in view, 50 nm. (B) Cryo-EM image of SMF-1 pili bundles sheared from *S. maltophilia*. Scale bar, 50 nm. The upper right inset shows a two-dimensional class average of a bundle comprising two SMF-1 pili. Pili bundles are indicated with green arrowheads. (C) The 4.0 Å cryo-EM reconstruction of an SMF-1 pili bundle (left), and the corresponding ribbon model built into the map (right). To clearly demonstrate the filament’s polarity, five different subunit conformations are colored distinctly, as indicated. The protein backbone of the atomic model is colored to match the map.

To directly determine the pilus identity, we sheared the pili from *S. maltophilia* via vortexing, and the soluble fraction was imaged using cryogenic electron microscopy (cryo-EM) to obtain an atomic pilus structure. With the advancement of cryo-EM^41^ and AI tools^42,43^, it is now routine practice in the field to identify protein components from a cryo-EM map better than 4 Å resolution. Under cryo-EM, most observed *S. maltophilia* pili appear in bundle forms with large diameters (Fig. 1B). This bundling-network morphology is not only seen in archaic CUP pili but has also been widely observed across various different pili and domains of life, where it has primarily been linked to biofilm formation. These observations, supported by experimental cryo-EM structures, include the Csu pilus^26,44^ (archaic CUP) of *Acinetobacter baumannii*, the CupE^25^ pilus (archaic CUP) of *Pseudomonas aeruginosa*, TasA pilus^19^ from *Bacillus subtilis*, archaeal bundling pili^20,21^ from *Pyrobaculum calidifontis* and *Pyrodictium abyssi*, and bacterial cytochrome nanowires^45^ in *Desulfuromonas soudanensis* WTL.

Among these other pili frameworks, it has been directly shown via cryo-EM that archaeal bundling pili^20,21^ and cytochrome nanowires^45^ pack in an antiparallel fashion. While the single filament structures of CupE^25^ and Csu^26^ pili were originally reported, a very recent study^44^ of Csu pili demonstrated that they pack as antiparallel filaments with several interacting modes. The TasA pili bundles, however, were suggested to possibly pack in both parallel and antiparallel fashions based on molecular dynamics simulations^19^, even though it is not clear how parallel-packed filaments would possibly contribute to biofilm formation. Here, in *Stenotrophomonas maltophilia*, the cryo-EM reconstruction reached 4 Å for a “pili-couple” structure containing two filaments (Fig. 1C). This resolution is sufficient for us to clearly identify the pilin as SMF-1 (BDLECG_02200, Fig. S1), rather than the other three proteins with a similar fold in the proteome (BDLECG_02926, BDLECG_03815, BDLECG_03816). The pili-couple packs in an antiparallel fashion, and notably, no parallel packing classes were found during 3D classification. Interestingly, from iterative 2D classification, the majority of classes consist of two pili where the subunits within both pili are clearly resolved. In classes containing three or more pili, the subunits are either not well-resolved or the density of those filaments is not at a similar level (Fig. S2). This suggests that in SMF-1 bundles, the interface between a pili-couple is very specific, while the interfaces become quite variable when these pili-couple units further bundle into a larger framework.

### Interactions between SMF-1 pilins that hold bundles together

Based on their diameters, many observed SMF-1 bundles (Fig. 1B) clearly contain more than two pili. Through cryo-EM, we reveal that these bundles are composed of “pili-couple” units consisting of two antiparallelly packed filaments. These units can further pack into wider bundles via non-specific interfaces. We also observed sparsely distributed single filaments under cryo-EM; despite a limited number of single-pilus particles, the cryo-EM reconstruction reached ∼6 Å resolution, yielding refined helical parameters of a 26.4 Å rise and −152.2° rotation. These helical parameters and pilin arrangements are almost identical to those found within the subunits of the pili-couple unit (Fig. S3). Interestingly, closer examination of the pili-couple unit suggested the existence of a global helical symmetry, where the pilin returns to an identical environment every five subunits (Fig. 2A). To test this, we re-extracted the particles into large boxes and performed helical refinement using five times the helical parameters of a single pilus. As expected, global helical symmetry was confirmed, and the helical-averaged volume reached 4.7 Å without any extra local refinement. Given the antiparallel arrangement, there is also a D1 symmetry present, though it is not easily identified by standard software (Fig. S3).

**Figure 2.**
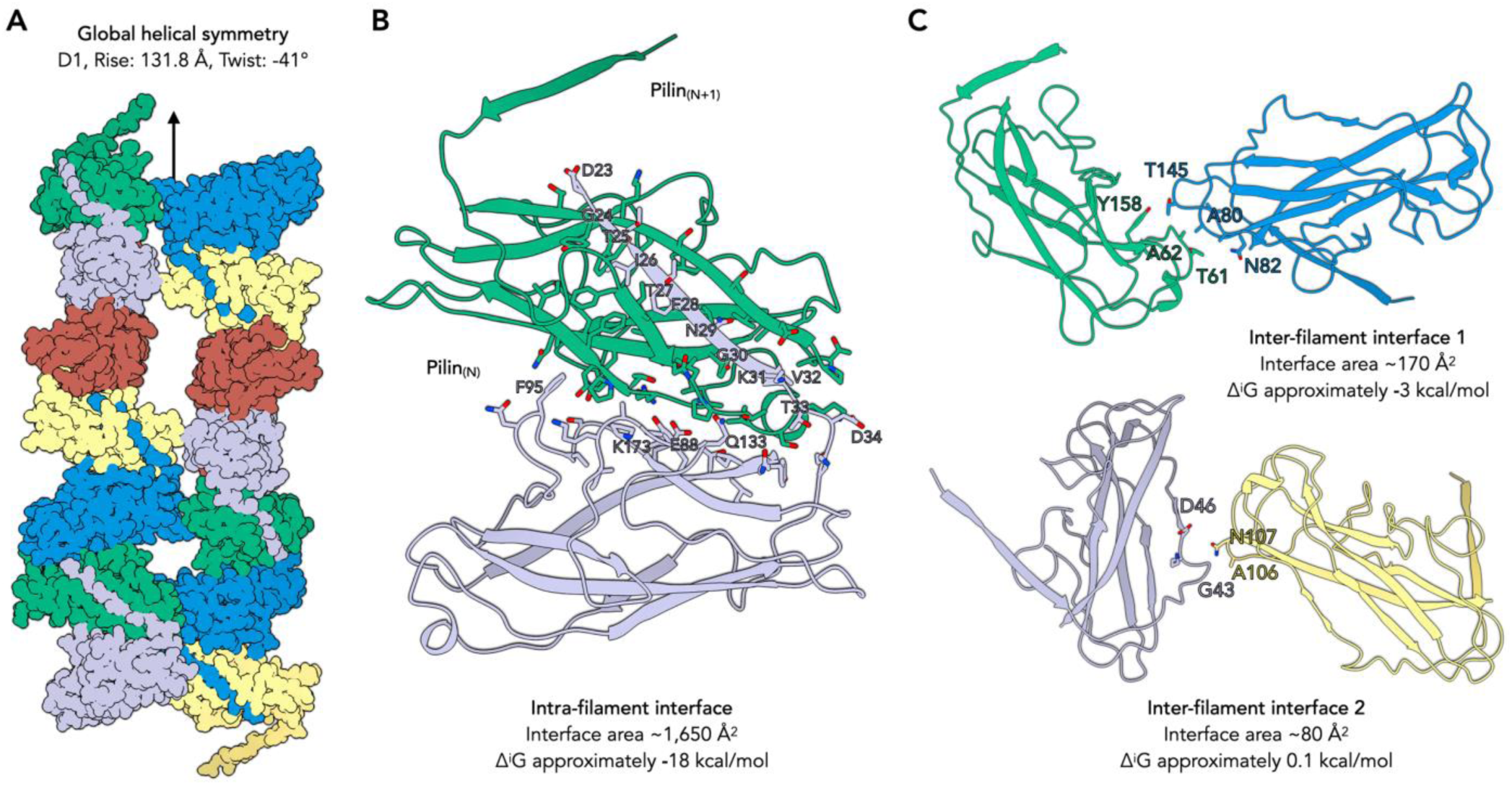
Pilin-pilin interfaces within the SMF-1 bundles. (A) Atomic model of SMF-1 pili-couple revealing a global helical symmetry within these two filaments, in which the asymmetrical unit is composed of 10 pilins. Pilins with identical environments are indicated by the same color, consistent with the color scheme used in Fig. 1. (B) Pilin-pilin interface within the same pilus. This interface is largely formed by the donor strand of one pilin inserting into its neighboring pilin subunit. Amino acids of the donor strand, as well as large side chains outside the donor strand that contribute to the interface, are labeled. (C) Pilin-pilin interface between different pili. Two types of interfaces are present: the larger interface is shown at the top, and the smaller interface is shown at the bottom.

Within a single pilus filament, a strong interface exists between pilin_(N)_ and pilin_(N+1)_. This interface is largely comprised of two components (Fig. 2B): (1) the donor strand of pilin_(N)_ inserting into pilin_(N+1)_ to form β-sheet hydrogen bonds and additional hydrophobic interactions, and (2) extensive hydrogen bonding at the interface of the two pilins’ major bodies. The total buried surface area of this interface is 1,650 Å^2^, which is typical for pilin-pilin interfaces and notably large for such a small protein. Regarding the detected inter-filament contacts, we observed two smaller unique interfaces mediated by hydrogen bonds, with interface area of 170 Å^2^ (Type 1) and 80 Å^2^ (Type 2), as estimated by PISA^46^ (Fig. 2C). While small, these interfaces accumulate rapidly along the bundles. For instance, the interface area between two adjacent SMF-1 filaments is 500 Å² per 13.2 nm. This scales to a substantial total interface of approximately 75,700 Å^2^ for two 2-µm long SMF-1 pili, a value that is further amplified in bundling arrays containing dozens of such pili-couple units.

### The archaic SMF-1 pilus was annotated as a classic CUP pilus

Despite its quaternary structure clearly identifying it as an archaic pilus, the SMF-1 pilus was historically annotated as a classic CUP pilus (γ clade)^31,47^. The classification of CUP pili is largely based on the fimbrial operon architecture as well as the sequence of the usher protein. When comparing the SMF-1 operon with other CUP operons of known cryo-EM structures, the SMF-1 operon shares similarities with both the classic type 1 pilus and two archaic CUP pili (Fig. 3A). All of these operons follow the same gene order: major pilin, chaperone, usher, and tip adhesin. However, the SMF-1 operon contains only these core genes, while other CUP systems include additional accessory genes (Fig. 3A). Based on seminal evolutionary analyses of CUP operons^47^, two clades could match the exact gene number and order of SMF-1: either the classic CUP pilus (γ clade) or the archaic CUP pilus (σ clade).

**Figure 3.**
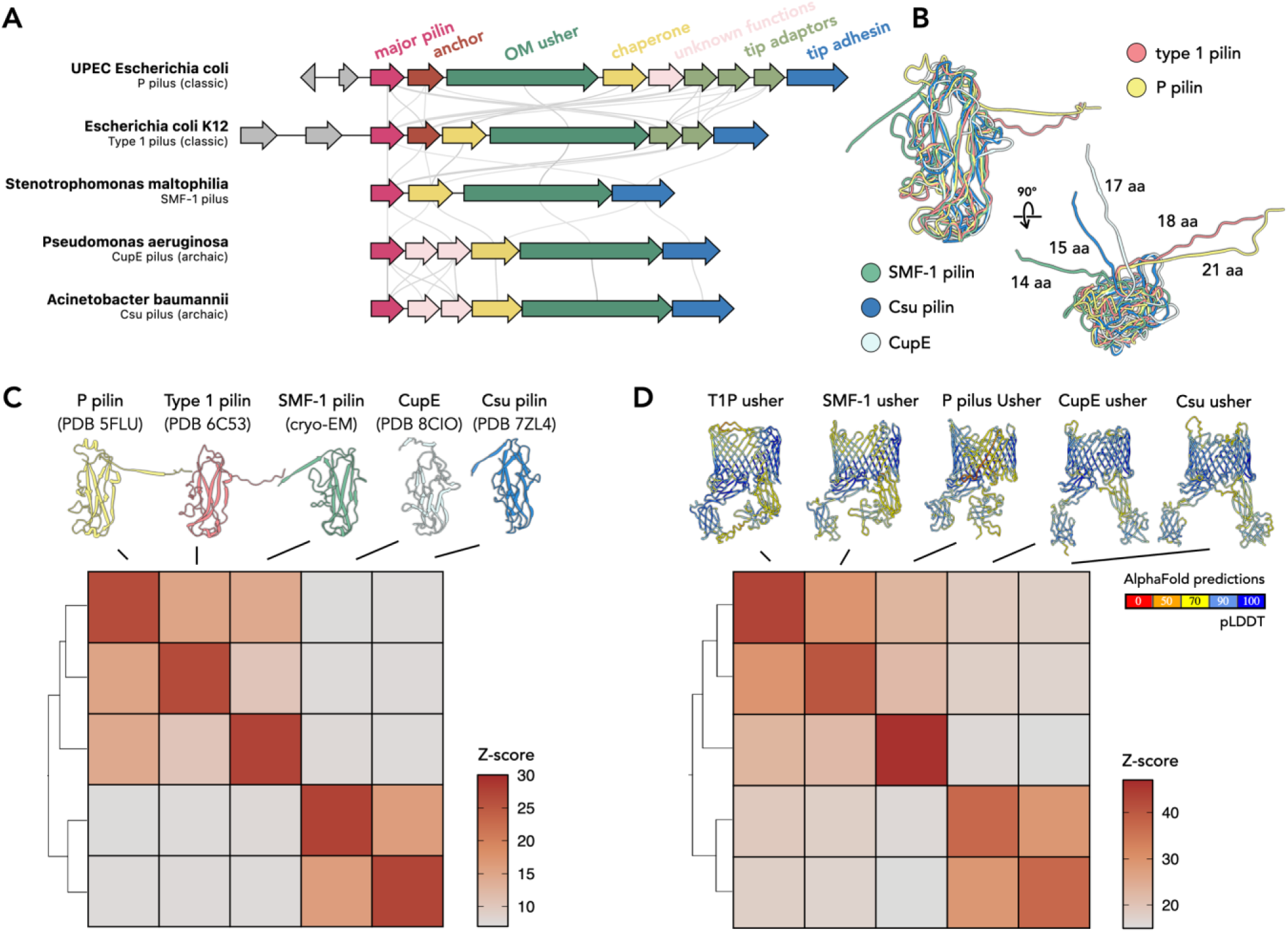
Comparison of the SMF-1 pilus with known classic and archaic CUP pili. (A) Gene structures of the SMF-1 pilus operon and other CUP pilus clusters identified in various species, including the P pilus (classic), Type 1 pilus (classic), CupE pilus (archaic), and Csu pilus (archaic). Genes with similar known functions are assigned the same color, with corresponding annotations indicated at the top. Gray lines link genes between clusters that share a sequence identity greater than 20%. (B) Structural alignment of five CUP pilins, with donor strand lengths labeled accordingly. To be consistent, the last residue of the donor strand was defined as the residue immediately before the conserved cysteine residue involved in disulfide bond formation. (C) DALI all-to-all analysis of the five CUP pilins. A structural similarity matrix is shown with an accompanying dendrogram on the left. Structural similarity is represented by DALI Z-scores, and the corresponding PDB codes used for the analysis are labeled. (D) DALI all-to-all analysis of five usher proteins. A structural similarity matrix is shown with an accompanying dendrogram on the left. These usher proteins were predicted using AlphaFold3; the displayed models are colored according to pLDDT confidence measurements.

Next, we examined the pilin structures of classic and archaic CUP pili determined by cryo-EM. Unsurprisingly, despite very low sequence identity between some of the pilins, their main bodies all possess an Ig-like fold and align well (Fig. 3B). However, the length of the donor strands and the angles at which they depart from the main body differ significantly. Archaic CUP pilins generally have shorter donor strands (14–17 amino acids), whereas donor strands in classic CUP pilins are longer (18–21 amino acids) and tilt at a reverse angle (Fig. 3B). The extra length of the donor strands in classic CUP pili likely indicates they form contacts between pilin_(N)_ and pilin_(N+2)_, which could further stabilize the rod-like tubular structure contacts of classic CUP pili.

Finally, to determine at the protein-fold level whether SMF-1 is closer to classic or archaic CUP pili, we compared the pilin structures (Fig. 3C) and AlphaFold-predicted usher structures (Fig. 3D) using a DALI^48^ all-to-all analysis. Surprisingly, in both cases, the folds of the SMF-1 pilin and usher appear much more similar to known classic CUP pili than to archaic ones. It is therefore unsurprising that the SMF-1 pilus was previously annotated as a classic rather than an archaic pilus. Our cryo-EM structure of SMF-1 reveals that such sequence-based classifications are not always accurate; instead, SMF-1 might represent an intermediate example of CUP pilus functional evolution, transitioning from a zigzag shape promoting biofilm formation to a rod-like shape optimized for robust adherence.

### SMF-1 pilin lacks the interface to form a rod-like classic CUP pilus

Since the sequences of the SMF-1 pilin and usher are remarkably similar to classic CUP pili, we investigated why SMF-1 does not form a rod-like structure similar to those systems. To explore this, we first performed a structure-based sequence alignment and manually validated that at every turn of the Ig-like fold, residues at corresponding positions across the five analyzed pilins aligned at the sequence level (Fig. 4A). In classic CUP pili, the key interfaces^23^ maintaining the rod-like architecture are the contacts between pilin_(N)_ and pilin_(N+3)_. Interestingly, these residues are not conserved between the two classic CUP pili (P pilus and type 1 pilus) and are quite different among all five analyzed pilins (Fig. 4A). Given that these are mostly hydrophilic residues, we initially questioned whether this interface was mediated by extensive polar contacts. However, examination of the electrostatic surfaces at the pilin_(N)_ and pilin_(N+3)_ interface ruled this out, as none of the five pilins presented strong polar surfaces (Fig. S4).

**Figure 4.**
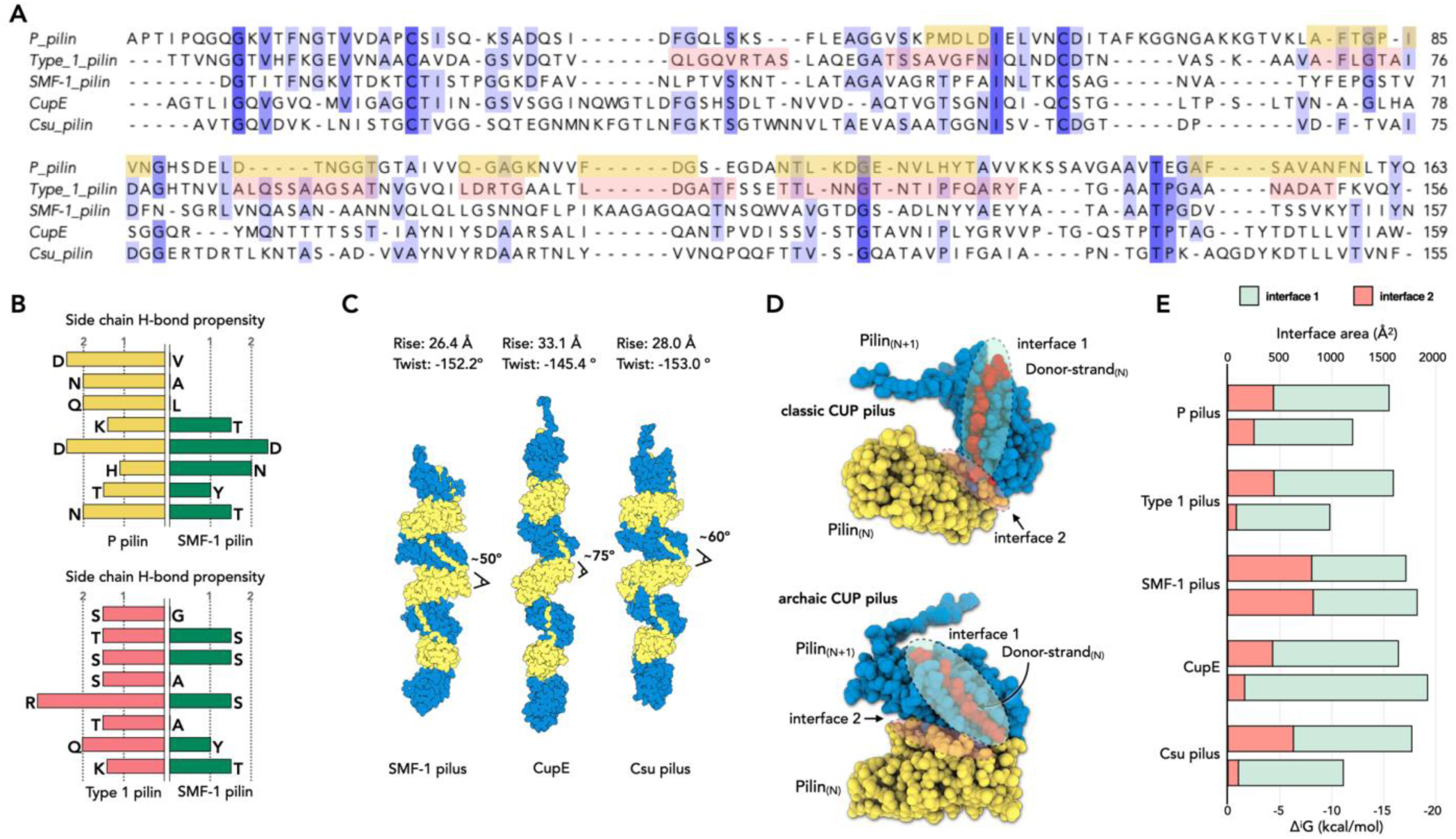
SMF-1 lacks the interface required to form a rod-like classic CUP pilus. (A) Structure-based sequence alignment of five CUP pilins. For the two classic CUP pili, residues involved in the interface between pilin_(N)_ and pilin_(N+3)_ are highlighted in yellow and red, respectively. (B) PDB-PISA analysis of the classic CUP pilus interface between pilin_(N)_ and pilin_(N+3)_. Several residues were identified forming hydrogen bonds with side chains that are more than 50% buried. The side-chain H-bond propensity of these residues is compared to the corresponding residues in SMF-1 based on the sequence alignment in (**A**). (C) Atomic model of a single SMF-1 pilus, shown in comparison with two archaic CUP pili: the CupE pilus and the Csu pilus. (D) Comparison of the interface between pilin_(N)_ and pilin_(N+1)_. The classic CUP pilus interface is shown at the top, while the archaic interface is shown at the bottom. Beyond “interface 1” formed by the donor strand of pilin_(N)_ and main body of pilin_(N+1)_, a second “interface 2” exists between the two subunits, which is largely mediated by hydrogen bonds. (E) A detailed comparison of the interfaces described in (**D**) across the five pili. For each pilus, two bars are provided: the top bar represents the interface area (Å²) and the bottom bar represents the estimated solvation free energy gain upon interface formation (kcal/mol). The relative contributions from interface 1 and interface 2 are distinguished by different colors.

A detailed analysis using PISA revealed that the pilin_(N)_ and pilin_(N+3)_ interface is largely stabilized by hydrogen bonds. Classic CUP pilins possess several residues with side chains that are more than 50% buried and contribute to hydrogen bonding. In contrast, the corresponding residues in the SMF-1 pilin have a lower propensity^49^ to form side-chain hydrogen bonds (Fig. 4B). Furthermore, all archaic CUP pilins contain a loop region located within the hypothetical pilin_(N)_ and pilin_(N+3)_ interface that would make the formation of such an interface energetically unfavorable.

Because archaic CUP pilins lack the pilin_(N)_ and pilin_(N+3)_ interface, we wondered whether their structures are more flexible compared to classic CUP pili. It has been previously reported that it takes approximately 25 pN to unwind classic CUP pili by breaking the pilin_(N)_ and pilin_(N+3)_ interface^22^. While it requires less force to extend and unwind archaic pili^26^, the unwinding phase itself still occurs at approximately 25 pN. This suggests that the primary pilin_(N)_ and pilin_(N+1)_ interface in archaic CUP pili is also quite strong. From a cryo-EM perspective, the fact that these archaic CUP pili can reach near-atomic resolution indicates that their contacts are consistent across different filaments, allowing them to be averaged and reconstructed at high resolution. Among the three known archaic pili structures, all share a helical rise of approximately 30 Å and a twist of around −150° (Fig. 4C). Notably, in SMF-1, the interface between pilin_(N)_ and pilin_(N+1)_ includes not only a large interface between the donor strand and the main body but also significant contacts between the two main bodies (Fig. 4D). In general, archaic CUP pilins appear to have larger contacts between pilin_(N)_ and pilin_(N+1)_ compared to classic CUP pili, which is consistent with this being the primary interaction holding the pilus together.

### SMF-1 pili production leads to biofilm initiation

While it has been reported that deletion of the *smf-1* gene leads to a dramatic reduction in biofilm formation across various systems^36^, it has also been recently appreciated that pili production is often a highly dynamic process^50,51^. To determine if biofilm formation has a direct correlation with the production of extracellular SMF-1 pili, and whether other appendages contribute to biofilm initiation, we grew *S. maltophilia* in three different liquid media at 37°C for 24 hours and examined their appendage production via negative staining. As a result, in BHI medium, almost all *S. maltophilia* cells remained “bald”, producing only short SMF-1 pili (Fig. 5A). Conversely, in CDM and M9CA (with glucose) media, much longer SMF-1 pili were observed (Fig. 5B-C). In M9CA (with glucose) medium, flagellar filaments are also spotted on some cells, whereas flagella production were not observed in CDM medium (Fig. 5B).

**Figure 5.**
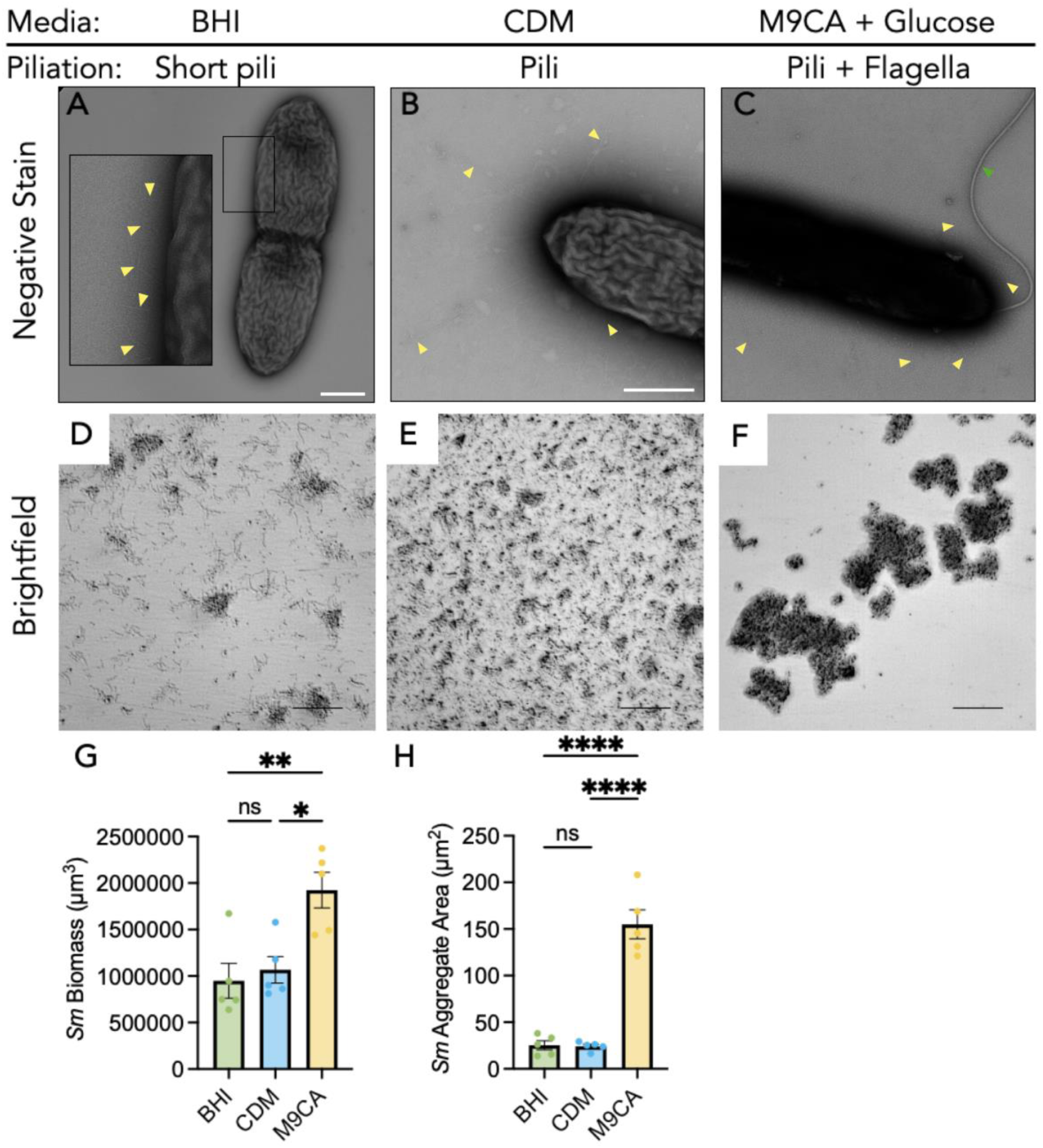
*S. maltophilia* forms aggregates in growth conditions that promote production of pili and flagella. (A-C) Negative staining TEM to show pilus and flagella production. SMF-1 pili are indicated with yellow arrowheads; Flagellum is indicated with green arrowhead. (D-F) Representative minimum intensity projections of image stacks obtained with brightfield microscopy show *S. maltophilia* adherence and aggregate formation in BHI (D), CDM (E) and M9CA (with glucose) (F) media following 24 hours of static culture. Scale bar, 80 µm. (G-H) Quantification of total *S. maltophilia* biomass (G) and aggregate area (H) in each growth condition. Data from 5 biological replicate experiments, with 5 fields of view quantified per condition per experiment. Welch’s t-test performed on biomass volume and aggregate area data (ns = nonsignificant, *P≤0.05,**P≤0.01, ****P≤0.0001). Data represents mean ± SEM.

We next evaluated bacterial aggregation in these specific growth conditions to determine the relationship of SMF-1 pili production to biofilm initiation. Brightfield microscopy identified differences in *S. maltophilia* aggregation between the three media (Fig. 5D-F). In BHI, where the bacteria lacked long pili, *S. maltophilia* exhibited a low level of aggregation, with many individual surface-attached cells visible. In contrast, numerous surface-attached aggregates were observed in CDM, where the bacteria produced longer pili. Despite these morphological differences, the total biofilm biomass (Fig. 5G) and aggregate size (Fig. 5H) were similar between the BHI and CDM conditions. Interestingly, *S. maltophilia* formed large aggregates in M9CA (with glucose) media, where bacteria produced numerous pili as well as some flagella (Fig. 5F). This is consistent with reports in other bacteria noting that flagellar production often promotes early biofilm initiation^52–56^. This growth condition resulted in significantly increased total biomass and aggregate size after 24 hours compared to the other media types (Fig. 5G,H), suggesting SMF-1 pili and flagella work in tandem to support robust aggregation.

## DISCUSSION

In this study, our cryo-EM structure reveals that SMF-1, previously annotated as a classic CUP pilus, is in fact an archaic CUP pilus that forms thick bundles between cells. This architecture provides a structural explanation for why deletion of *smf-1* leads to a dramatic reduction in biofilm formation^36^. Specifically, we captured the building units of these bundles, which consist of two SMF-1 filaments packed in an antiparallel arrangement. Repeated assembly of dozens of such units would generate a robust tethering platform that mechanically links different cells together via their pili. We further connected these extracellular appendages to biofilm initiation by monitoring *S. maltophilia* pilus production and aggregate formation under defined growth conditions; we found that SMF-1 pili were required for formation of large cellular aggregates, a phenotype likely relevant to persistence and chronic infections, and we found that flagella together with pili can further accelerate this process.

Why would bundled pili facilitate colonization and biofilm formation? If filaments within these bundles originate from different cells and are packed antiparallel, the functional consequence is straightforward: the bundle provides strong, multivalent intercellular connections. This phenomenon appears to be widespread among microbial extracellular appendages and may represent a general strategy for early colonization and biofilm initiation. To date, among examples for which multi-filament cryo-EM structures are available, including archaic CUP pili^44^, cytochrome nanowires^45^, and archaeal bundling pili^20,21^, all bundles are consistently antiparallel-packed. Moreover, even in cases where an atomic model is not available but 2D averages are clear (for example, bacterial spore appendages), bundles likewise appear to form through antiparallel organization^57^. This does not rule out the possibility of parallel-packed pili bundles; indeed, multiple pili from the same cell could pack together transiently. However, it seems mechanically unlikely that an ordered, parallel-packed arrangement would make a major contribution to intercellular tethering/colonization in the relatively unconstrained extracellular space. By contrast, in the more complex environments of eukaryotic cells, filament bundling (e.g., actin) supports diverse functions, and both parallel and antiparallel orientations are observed^58^.

Comparison of archaic and classic CUP pili operons (Fig. 3A) highlights another notable distinction: the three known archaic systems lack an anchor pilin that terminates assembly. In some classic CUP pili, termination occurs when an anchor pilin binds the chaperone with high affinity, preventing chaperone dissociation and leaving the donor-strand binding groove occupied and unable to accept additional major pilins^59^. No analogous anchor pilins have been identified in archaic CUP systems. Consistent with the remarkably simple gene cluster of SMF-1, we likewise do not identify a clear candidate protein to fulfill this role. This raises the question of whether archaic CUP pili truly require an anchor pilin for termination. One possibility is that the termination is an emergent property of bundling. As dozens of SMF-1 pili filaments assemble into a bundle, inter-filament contacts accumulate over long distances. Extending a filament within an established bundle would require disrupting many of these interfaces to accommodate insertion of a new subunit, potentially making continued growth energetically unfavorable. In this scenario, a dedicated anchoring mechanism may be less critical than in classic CUP pili, which primarily function as shock absorbers and may require a strong connection to the outer-membrane usher.

From an evolutionary perspective, it is intriguing that SMF-1 pilin and usher sequences and their overall structures closely resemble those of classic CUP pili, whereas the supramolecular architecture is clearly archaic. The γ_4_ sub-clade (where SMF-1 is classified) may represent an evolutionary intermediate, assuming all CUP pili share a common ancestor. Alternatively, the distinction between classic and archaic CUP pili may be less absolute than current classifications imply; some classic CUP pili might adopt a zigzag, archaic organization under particular mechanical forces or environmental conditions. Future studies that systemically probe how sequence features, assembly constraints, and environmental cues shape CUP pilus architecture will be essential for clarifying their functional diversity and evolution relationships.

## METHODS

### *S. maltophilia* SMF-1 pili preparation for cryo-EM

*S. maltophilia* cells were grown aerobically at 37 °C for 48 hours in Tryptic Soy Broth (TSB, BD™) medium within a 5% CO_2_ incubator, as previously reported^40^. The resulting pellet was resuspended in 4 mL of PBS buffer, and the cell suspension was processed in a homogenizer for 10 min to shear off the extracellular filaments. Following homogenization, the cells were removed by centrifugation at 10,000 × *g* for 10 min. The supernatant was then collected, and the filaments were pelleted via ultracentrifugation (Beckman 50.3 Ti rotor, 108,000 x *g*, 1.5 h, 4°C). After the run, the supernatant was discarded, and the pellet was resuspended in 150 µL of PBS. Finally, DNase I (NEB) was added to the sample to remove any remaining extracellular DNA fibers.

### Cryo-EM conditions and image processing

The cell appendage sample (ca. 4.5 μl) was applied to glow-discharged lacey carbon grids and then plunge frozen using an EM GP2 Plunge Freezer (Leica). The cryo-EM micrographs were collected on a 300 keV Titan Krios with a K3 camera at 1.11 Å/pixel and a total dose of ca. 50 e/Å^2^. First, patch motion correction and CTF estimation were done in cryoSPARC.^60–62^ Next, particles were auto-picked by “Filament Tracer” with a shift of approximately 28 Å. All auto-picked particles were extracted into a 320-pixel box and subjected to multiple rounds of 2D classification. To clearly reveal how SMF-1 pili are packed within the bundles, all classes containing two filaments (totaling 85,928 particles) with apparent separation of single pilins were selected for 3D reconstruction. Because this reconstruction includes two filaments, it was initially unclear whether a global helical symmetry existed; therefore, we began with Ab-Initio reconstruction. After visual inspection of the resulting volume, we further improved the resolution through successive rounds of “Homo Refine”, “Local CTF Refinement”, a second “Homo Refine”, and a final “Local Refine”. This reconstruction reached a resolution of 4.0 Å, with beta-sheets well separated and amino acid side chains clearly resolved.

The hand of the 3D volume was determined by comparison with the predicted pilin structure and existing archaic CUP pilin proteins. The helical symmetry of the individual filament was determined from the built atomic model, exhibiting C1 symmetry, a helical rise of 26.4 Å, and a twist of −152°. From the model, it is also evident that the pili-couple structure has a global D1 helical symmetry. In this configuration, five copies of pilin from one filament serve as the asymmetrical unit, with helical parameters five times those of the pilin within a single filament (rise 131.8 Å, twist −41°). This observation was validated by applying the global symmetry to the pili-couple particles in a reconstruction using a larger 448-pixel box. The resolution of the final reconstruction was estimated using Map-to-Map and Model-to-Map FSC. The final volume was then sharpened using EMReady^63^, and the statistics are summarized in Supplementary Table 1.

### Model building of SMF-1 pilus

The first step of model building involved identifying the pilin protein from the experimental cryo-EM map. Among all proteins with a similar fold (*BDLECG_02200*, *BDLECG_02926*, *BDLECG_03815*, *BDLECG_03816*) in this strain, the pilin protein produced by gene *BDLECG_02200* was identified as the best fit. This identification was achieved using ModelAngelo^42^ combined with visual inspection of the model fit within the cryo-EM map. The AlphaFold^64^-predicted model of the identified pilin served as the starting point for the building process. A single pilin subunit was first docked into the map using ChimeraX^65^ and then manually refined in Coot^66^. Subsequently, a pili-couple model containing 14 pilin copies was generated and underwent real-space refinement in PHENIX without NCS (non-crystallographic symmetry) constraints^67^. MolProbity^68^ was used to evaluate the quality of the resulting filament model. The refinement statistics for both the mono-pili and tri-pili are provided in Supplementary Table 1. Data visualization was primarily performed in ChimeraX^65^.

### Negative staining

Continuous carbon film grids (Electron Microscopy Sciences, 300-mesh, Cu) were used for negative staining. Briefly, the grids were glow-discharged for 30 s at 25 mA prior to sample application. Following this, 4 μL of liquid culture containing *S. maltophilia* cells and 10% DNase I was applied directly to the carbon side of the glow-discharged grid and incubated for 20 s. The grid was subsequently back-blotted with filter paper (Whatman) and gently dragged downward against the paper until the excess sample was removed. Immediately following this, the grid was washed twice by dropping 12 μL of ultrapure water onto the grid surface and quickly back-blotted as before. After the washing steps, 10 μL of 2% uranyl acetate (EMS) was applied to the grid and back-blotted. The grid was then allowed to air-dry for 1 min. Negative-staining TEM images were acquired using a JEOL JEM-1400 transmission electron microscope equipped with an AMT NanoSprint43 Mk-II camera at the University of Alabama at Birmingham’s High Resolution Imaging Facility. Pilus production was assessed across five cells selected at random positions from the grid.

### Monitoring *S. maltophilia* pili production and aggregation assay

Stocks of *Stenotrophomonas maltophilia* cells were streaked onto BHI (BD Diagnostics) agar plates and incubated in a humidified environment at 37°C for 72 hours. For the aggregation assay, a single colony was selected and inoculated into 5 mL of liquid culture medium; these cultures were then grown statically for 24 hours at 37°C with. Based on negative-staining results of pilus production, the following media were used for the aggregation assay: (1) BHI medium for non-piliated growth; (2) CDM^69^ for SMF-1 piliated growth; and (3) M9CA^70^ with glucose for both SMF-1 piliated and flagellated growth.

A single colony of *S. maltophilia* was inoculated into 5 mL of the specified liquid culture media and gently vortexed before transferring 200 µL to a sterile 96-well, non-coated flat bottom plate (Griener). Bacteria were further diluted 1:10 and statically incubated at 37°C for 24 hours. Images were taken on a Nikon Eclipse Ti2 widefield microscope (Nikon). At least 5 individual fields of view were compared per well from 5 independent experiments. Analysis of average aggregate size was conducted in the Nikon NIS-Elements AR software package (Version 5.42.02 Build 1801) as previously described^71^. Data analysis was performed in Microsoft Excel Version 16.105.2^72^. Welch’s t-tests were performed on biomass and aggregate size. *P* values were considered significant if less than or equal to 0.05.

## Data availability

The three-dimensional reconstruction of CT pilus has been deposited in the Electron Microscopy Data Bank with accession code EMD-75440. The atomic model has been deposited in the Protein Data Bank with accession code 10SS. The genome of *Stenotrophomonas maltophilia* strain was sequenced and deposited in NCBI BioSample database with accession code SAMN55362942.

## Acknowledgments

This research was, in part, supported by the National Cancer Institute’s National Cryo-EM Facility at the Frederick National Laboratory for Cancer Research under contract 75N91019D00024. Electron microscopy screening was carried out in the UAB Cryo-EM Facility, supported by the Institutional Research Core Program and O’Neal Comprehensive Cancer Center (NIH grant P30 CA013148), with additional funding from NIH grant S10 OD024978. Research reported in this publication was supported by the UAB High Resolution Imaging Facility. We are grateful to Dr. James Kizziah and Dr. Jenny Wang for assisting with the screening or data collection. The work in F.W. laboratory was supported by NIH grant GM159704. The work in M.K. laboratory was supported by Cystic Fibrosis Foundation grant KIEDRO24P0 and CFF grant DAVIS24R0 (Predoctoral training award) to C.S.

## Author contributions

H.Z. and J.L.F. performed pili sample preparation. J.L.F. performed the negative staining analysis. C.C.S., under the supervision of M.R.K., performed aggregation assays. H. Z., under the supervision of H.W. performed bacterial genome sequencing. J.L.F., A.Z., A.N.R., under the supervision of F.W. performed cryo-EM analysis and model building. F.W. M.R.K. and H.W. obtained funding. F.W. wrote the manuscript with input from all authors.

